# Intrinsic Network Activity Reflects the Fluctuating Experience of Tonic Pain

**DOI:** 10.1101/2021.06.30.450591

**Authors:** Bettina Deak, Thomas Eggert, Astrid Mayr, Anne Stankewitz, Filipp Filippopulos, Pauline Jahn, Viktor Witkovsky, Andreas Straube, Enrico Schulz

## Abstract

Although we know sensation is continuous, research on long-lasting and continuously changing stimuli is scarce and the dynamic nature of ongoing cortical processing is largely neglected.

In a longitudinal study with 152 fMRI sessions, participants were asked to continuously rate the intensity of applied tonic heat pain for 20 minutes. Using group independent component analysis and dual-regression, we extracted the subjects’ time courses of intrinsic network activity. The relationship between the dynamic fluctuation of network activity with the varying time courses of three pain processing entities was computed: pain intensity, the direction of pain intensity changes and temperature.

We were able to dissociate the spatio-temporal patterns of objective (temperature) and subjective (pain intensity/changes of pain intensity) aspects of pain processing in the human brain. We found two somatosensory networks with distinct functions: one network which encodes the small fluctuations in temperature and consists mainly of bilateral SI. A second right-lateralised network that encodes the intensity of the subjective experience of pain consists of SI, SII, the PCC, and the thalamus.

We revealed the somatosensory dynamics that build up towards a current subjective percept of pain. The timing suggests a cascade of subsequent processing steps towards the current pain percept.

## Introduction

Although we know sensation is continuous (James 1890; Fekete et al. 2018; Antony et al. 2021), laboratory research has largely focused on the processing of short, discrete stimuli. Research on long-lasting and continuously changing stimuli is scarce. The experience of tonic pain typically lasts longer than the more frequently investigated brief laser pain, whose cortical response is suggested to reflect the salience aspects rather than the pain (Legrain et al. 2011).

### Coding the intensity of long-lasting pain

The experience and intensity of tonic and chronic pain can fluctuate substantially over time (Mun et al. 2019) (Baliki et al. 2006; May et al. 2018). A number of studies (Lorenz et al. 2003; Pogatzki-Zahn et al. 2010; Favilla et al. 2014; Segerdahl et al. 2015; Nickel, May, Tiemann, Schmidt, et al. 2017; Davis et al. 2020) have specifically addressed the fluctuation of prolonged pain in healthy subjects, however, due to the inherent limitations of the application of pain (cutaneous and intramuscular saline injection, as well as incisions), most of these studies could not control the intensity of perceived pain. Consequently, they could not take habituation or sensitisation phenomena into account. As a result, the pain ratings gradually changed over time and appeared to reflect habituation (Favilla et al. 2014) or sensitisation (Nickel, May, Tiemann, Schmidt, et al. 2017). Likewise, experimental findings on cortical processes can obtain results that differ between the first and second half of the experiment (May et al. 2018).

In fMRI studies these low-frequency aspects of perception need to be controlled for as they are inevitably lost from the cortical data due to the required high-pass filtering of functional imaging sequences. However, a previous EEG study has shown that a continuous adaptation of the stimulus is feasible. The authors kept the subjective pain at a similar level and found gamma oscillations recorded at frontocentral electrodes to encode the subjective intensity of tonic pain (Schulz et al. 2015). Likewise, a further imaging study utilising arterial spin labelling (ASL) found the dorsal posterior insula to correlate with intensity of pain (Segerdahl et al. 2015).

### intrinsic network activity in pain research

In recent years neuroimaging studies on pain have largely shifted their focus to the analysis of intrinsic network activity. These studies have analysed the spatial characteristics of pain patients’ cortical maps in comparison to healthy control subjects and detected functional networks that are altered in chronic pain patients (Baliki et al. 2014; Androulakis et al. 2017). However, analyses on such network maps can not fully take into account the ongoing dynamics of the functional network. Furthermore, it is difficult to interpret how differences in single voxels would represent differences of the entire functional network.

Here, we aimed to investigate the temporal dynamics of intrinsic networks by relating their varying time courses to the time course of the individual fluctuations of the intensity of long-lasting painful heat. Consequently, we set a particular focus on the dynamic aspects of the intrinsic networks. In order to prevent a confound through sensitisation, we utilised a stimulation paradigm that inflicts perception-controlled tonic heat pain. In repeated recordings for each participant, we aimed to disentangle the cortical underpinnings of three entities of tonic heat processing: pain intensity encoding, temperature encoding, as well as the perception of rising and falling pain.

## Materials and Methods

### Participants

A group of 38 healthy subjects (18 female/20 male; aged 28 ± 5 years) was included in this study. All participants gave written informed consent. The study was approved by the Ethics Committee of the Medical Department of the Ludwig-Maximilians-Universität München (project number 19-756) and conducted in conformity with the Declaration of Helsinki. The subjects did not report any psychiatric, neurological or severe internal medical diseases. They did not take any medication and did not report any pain on the day of the MRI recording. Subjects with chronic or acute pain conditions or contraindications for an MRI examination (e.g. metal implants) were excluded from the study. The participants were remunerated 120€ for their contribution. All participants were recorded four times with a minimum gap of 2 days between sessions. A pretest outside the MRI on a day before the first recordings familiarised the participants with the stimulation and the rating procedure.

### Experimental procedure

During the recording of fMRI, heat pain was administered for 20 min to the upper part of the left forearm with a thermode (QST.lab; France). Participants rated the intensity of their ongoing pain using their right hand with an MRI-compatible potentiometer slider (Schulz et al. 2019). According to four different predefined and balanced time courses of pain intensity, the applied stimulus temperature was adapted continuously using a Matlab-based software controller (Figure 2). The design allows the pain perception to be kept at a predefined level. It thus controls for cortical habituation/sensitisation, variable thermodynamics of the underlying tissue, and peripheral (adaptation of peripheral nerves) processes that would otherwise lead to unbearable pain (at the end of the experiment) or no pain at all (at the beginning of the experiment). The applied temperature was limited to a maximum of 47 °C. None of the participants was suffering any harm, just a faint redness on the skin lasting a few hours.

**Figure1.**
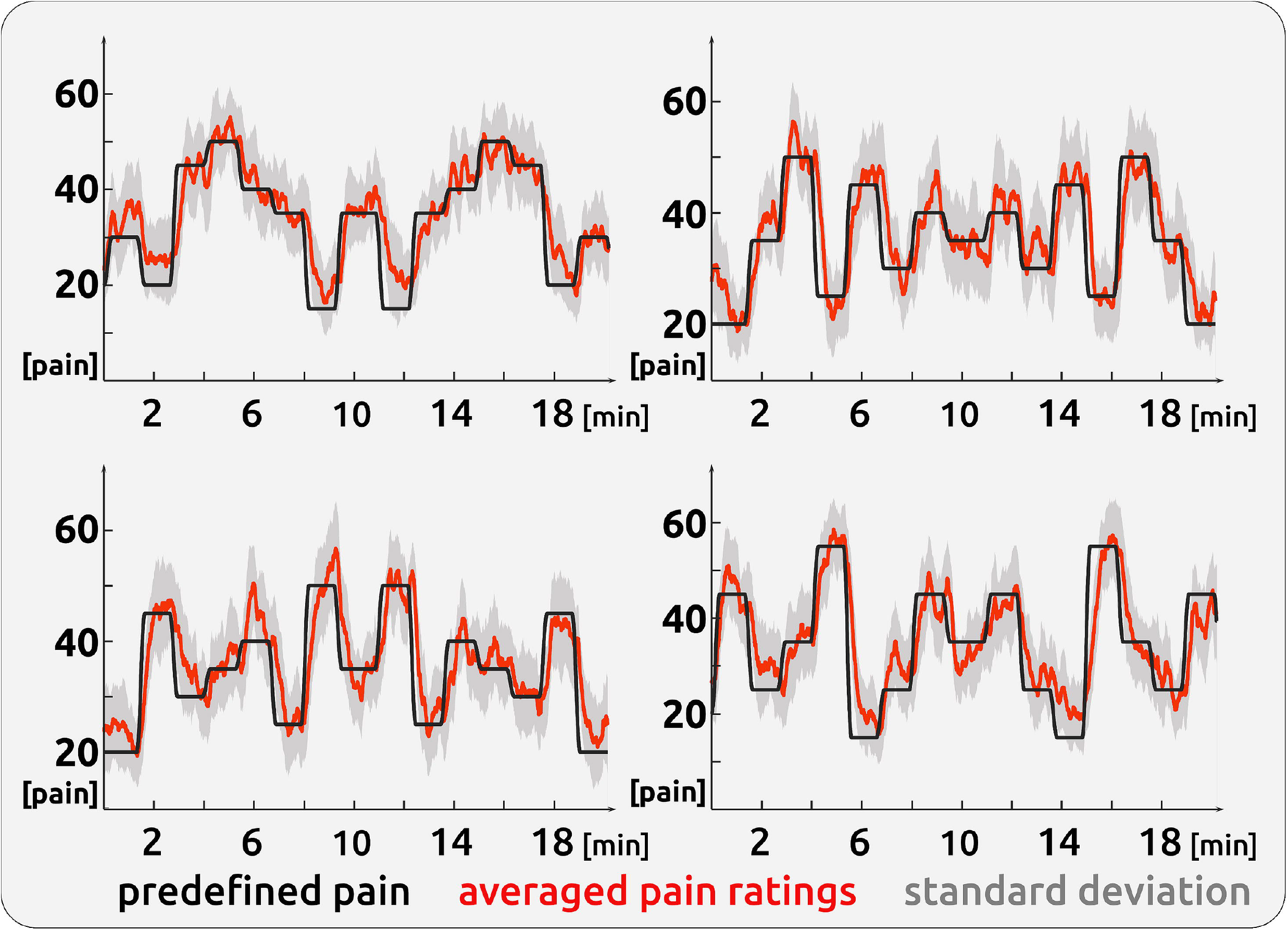
Time course of painful stimulation. The figure depicts the pain time courses of the 4 sessions that were to be experienced by the participants. The controlled application of heat confirmed the success of the software algorithm; the participants largely experienced and rated the inflicted pain according to the predefined time course. The standard deviation reflects the often unpredictable dynamics of subjective pain perception.

**Figure 2.**
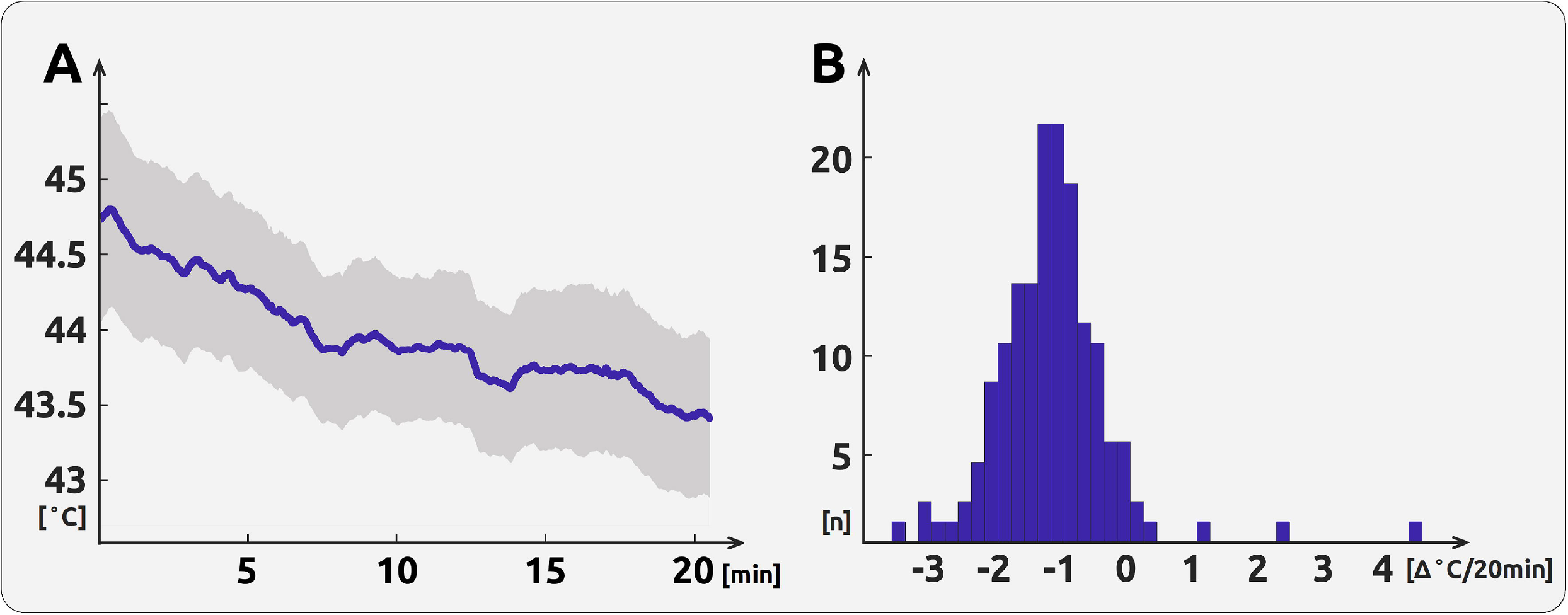
Time course of the pain ratings. The left figure (A) shows the averaged time course of thermode temperature across the 20 min of painful heat pain stimulation. The blue line represents the mean for each time point; the grey area is the standard deviation. The histogram on the right (B) confirms a habituation for only 7 out of 152 sessions. For most of the sessions (n) the participants sensitise.

The pain scale ranged from zero to 100 in steps of five with zero representing no pain and 100 representing the highest experienced pain. A red cursor on a dark grey bar (visual analogue scale, VAS) and a number above (numeric analogue scale, NAS) were shown on a screen during the entire experiment. The screen was visible through a mirror mounted on top of the MRI head coil. The intensity and the changes in perceived pain had to be indicated as quickly and accurately as possible. To minimise head movement, foams were placed around the head and participants were instructed to lie as still as possible.

### Data Acquisition

The data from 152 sessions (38 subjects x 4 sessions each) were recorded on a 3 tesla MRI scanner (Siemens Magnetom Skyra, Germany) with a 64-channel head coil. Using a multiband sequence (factor 4, T2*-weighted BOLD (blood oxygenation level dependent) images were acquired with the following parameters: number of slices = 44; repetition time/echo time = 760/30 ms; flip angle = 50°; slice thickness = 3 mm; voxel size = 3×3×3 mm^3^; field of view = 1470 mm. 1625 volumes were recorded in 1235 seconds. Field maps were acquired in each session to control for B0-effects. For each participant, T1-and T2-weighted anatomical MRI images were acquired using the following parameters for T1: repetition time/echo time = 2060/2.17 ms; flip angle = 12°; number of slices = 256; slice thickness = 0.75 mm; field of view = 240 mm, and for T2: repetition time/echo time = 3200/560 ms; number of slices = 256; slice thickness = 0.75 mm; field of view = 240 mm.

### Data processing - behavioural data

The rating data were continuously recorded with a variable sampling rate and down-sampled offline at 5 Hz. We applied the same filter to the rating and temperature data as to the imaging data (see below). For the statistical analysis, the resulting filtered time course was transferred to Matlab (Mathworks, USA; version R2018a) and down-sampled to the sampling frequency of the imaging data (1/0.76 Hz).

To disentangle the distinct aspects of pain intensity (AMP - amplitude) from cortical processes related to the sensing of rising and falling pain, we generated a further vector by computing the ongoing change (SLP - slope, encoded as 1, −1, and 0) in the pain ratings. The change is represented as the positive or negative slope of the regression of the least-squares line across a 3 s time window of the 5 Hz pain rating data. A vector of the absolute slope of pain ratings (aSLP - absolute slope, encoded as 0 and 1), represents periods of motor activity (slider movement), changes of visual input (each slider movement changes the screen), and decision-making (each slider movement prerequisites a decision to move).

The temperature vector (TEMP) was implicitly detrended by the filtering and represents the smaller high-frequency temperature changes. The 5 Hz data of AMP, SLP, aSLP, and TEMP vectors were convolved with a haemodynamic response function (HRF) implemented in SPM12 (Penny et al. 2011) with the following parameters: HRF = spm_hrf(0.2,[6 16 1 1 100 0 32]).

We shifted the 4 vectors between −25 s and 35 s in steps of 0.5 s (121 steps). These systematic shifts would account for:

a. the unknown delay of the BOLD response.
b. the unknown timing of cortical processing in reference to the rating: some ongoing cortical processes may influence later changes in pain ratings, other processes are directly related to the rating behaviour, or are influenced by the rating process and are occurring afterwards.
c. the unknown duration of the hemodynamic response and its variability.

During the process of vector generation the behavioural data were downsampled to the sampling rate of the imaging data. We are aware that the variable timing of the BOLD response and the variable timing of the cortical processes are intermingled and would interpret the timing aspects with utmost caution.

### Data processing - imaging data

Functional MRI data were preprocessed using FSL (Version 6.0.4, (Jenkinson et al. 2012), which included removal of non-brain tissue (using brain extraction, BET), slice timing correction, head motion correction, B0 unwarping, spatial smoothing using a Gaussian kernel of FWHM (full width at half maximum) 6 mm, and linear and non-linear registration to the Montreal Neurological Institute (MNI) standard template. High-pass temporal filtering was applied; the relatively long cutoff of 400 s was required due to the low-frequency changes of pain stimulation.

The data were further cleaned of artefacts by performing single-session independent component analyses (ICA) with MELODIC (Salimi-Khorshidi et al. 2014). Artefact-related components were evaluated according to their spatial or temporal characteristics and were removed from the data (Kelly et al. 2010; Griffanti et al. 2014). The average number of removed artefact components was 26 (±5). We deliberately did not include any correction for autocorrelation, neither for the processing of the imaging data nor for the processing of the pain rating time course, as this step can potentially alter the natural evolution of the processes we aim to investigate (see correction for multiple testing below). Head movement was uncorrelated to any of the behavioural vectors.

### Statistical analysis - imaging data

In a next step, we ran a group ICA with temporally concatenated data of all 152 recordings using MELODIC on a high-memory cloud server by disabling the “MIGP” option. The number of components was restricted to a reasonable number (100). In order to derive the fluctuating network time course for all components and for each of the sessions, dual regression was used (Nickerson et al. 2017). As a major advantage, the combination of ICA and dual regression can disentangle the involvement of the same region in different networks. At the same time, this increases the SNR, one the one hand, as distinct cortical signals are not overlapping and, on the other hand, noisy aspects of the data are separated as they are forming their own component. Please note that although we were using ICAs to determine intrinsic networks, this is not a study on resting-states.

Using Linear Mixed Effects models (LME; MixedModels.jl package in Julia; (Bezanson et al. 2015), we aimed to determine the relationship between fluctuating pain intensity and the fluctuating cortical activity separately for each component. The fluctuating network activity of a particular component was modelled through the time course of the four variables (AMP, SLP, aSLP, TEMP).

The statistical model is expressed in Wilkinson notation; the included fixed effects (network ~ AMP + SLP + aSLP + TEMP) describe the magnitudes of the population common intercept and the population common slopes for the relationship between cortical data and pain perception. The added random effects (e.g. AMP - 1 | session) model the specific intercept differences for each recording session (e.g. session specific differences in pain levels or echo-planar image signal intensities):

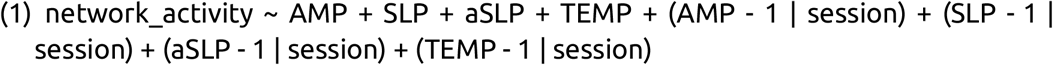

Each model was computed 121 times along the time shift of the behavioural vectors (−25 to 35 s in steps of 0.5 s, see above).

### Statistical analysis - correcting for multiple testing

All statistical tests were corrected for multiple testing (components, time shifts) and autocorrelation in the behavioural data: we created 1000 surrogate time courses using the IAAFT algorithm (Iterative Amplitude Adjusted Fourier Transform) from the original rating data, which were uncorrelated to the original rating data but had the same autocorrelation structure as the original data (Schreiber and Schmitz 1996). Using surrogate data, the entire LME analysis, including the temporal shifts, was repeated 1000 times, resulting in 1000*121*100 statistical tests for AMP, SLP, aSLP and TEMP. The highest absolute t-values of each repetition across all components and shifts were extracted. This procedure resulted in a right-skewed distribution of 1000 values for each condition. Based on the distributions of 1000 values (for AMP, SLP, aSLP, and TEMP), the statistical thresholds were determined using the “palm_datapval.m” function publicly available in PALM (Winkler et al. 2014, 2016).

## Results

### Behavioural data

In order to keep the subjective pain at a predefined level, the applied heat needed to be continuously controlled. The averaged thermode temperature was 44 °C (±0.5 °C, mean and standard deviation across the 152 sessions’ 20 min average), which dropped on average by 1.13 °C (±0.88 °C, computed as linear regression/20 min) across the time course of the 20 min experiment. The average of the applied thermode temperature was 44.7 °C in the first minute (mean across the 152 sessions’ first minute average) and 43.4 °C in the last minute (mean across the 152 sessions’ last min average). The applied mean temperature for a participant varied on average 0.56 °C (±0.36; max/2-min/2) across the 4 sessions.

### Imaging data

By using an LME, we were able to disentangle the different cortical underpinnings of (a) pain intensity (AMP), (b) the direction of the change of pain intensity (SLP), (c) temperature (TEMP), and (d) task-related decision-making, visual processing, and motor response (aSLP). The latter is not relevant for the present study; the results are therefore focussed on AMP, SLP, and TEMP.

### AMP - pain intensity encoding

We found a number of cortical networks whose time course exhibited significantly positive relationships with the time course of the subjective intensity of tonic pain (AMP; Figure 3, Table 1). The intensity of tonic pain was related to the activity of a large right-lateralised network consisting of inferior frontal, temporal and marginal regions (#29), and a right-lateralised somatosensory network that includes the primary (SI) and secondary (SII) somatosensory cortices, the anterior part of the posterior cingulate cortex (PCC), the posterior part of the anterior cingulate cortex (ACC), as well as the thalamus (#1). A third positively-correlated bilateral network consists of the paracingulate gyrus, the precuneous and the orbitofrontal cortex (#20), and a fourth network includes the bilateral medial orbitofrontal cortex and the left posterior part of the anterior cingulate cortex (ACC, #10). Further positively-correlated areas involve prefrontal regions that are connected to the cerebellum (#69), the insular cortex (#34), and the occipital cortex (#22). In contrast, there are several networks that exhibited decreased activity, predominantly in the occipital cortex (#66, #0), in prefrontal areas (#94, #72, #78), and in the cerebellum (#14, Table 1).

**Figure 3.**
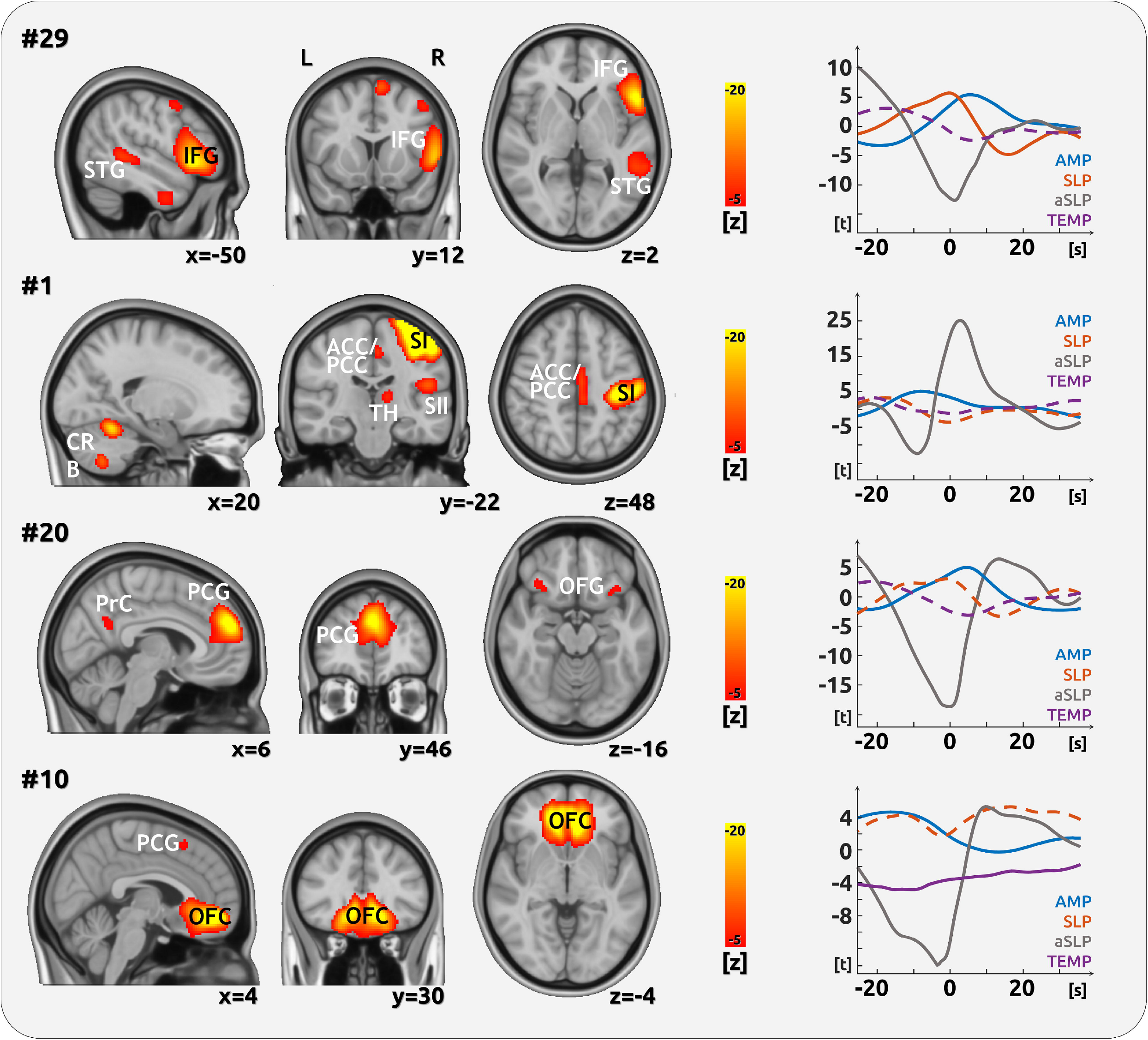
Encoding of the amplitude of pain intensity ratings. The figure shows intrinsic functional networks that encode the intensity (AMP) of tonic pain. The left side depicts the spatial characteristics of the networks; they were defined by the ICA. The significance of the contribution of each voxel to the network is represented by z-values. Only the right side depicts pain encoding. The graph shows the timing of the network activity in reference to the current pain rating at time point 0s. Solid lines indicate significant relationships, dashed lines indicate non-significant relationships. Please note the limited interpretability of the timing aspects due to the unknown lag of the haemodynamic response function, which may be different for each network. However, there is a *relative* delay for the change encoding compared to the pain intensity encoding for component #29. (Abbreviations: STG superior temporal gyrus, IFG inferior frontal gyrus, CRB cerebellum, ACC anterior cingulate cortex, PCC posterior cingulate cortex, SI primary somatosensory cortex, SII secondary somatosensory cortex, TH thalamus, PrC precuneous, PCG paracingulate gyrus, OFC medial orbitofrontal cortex).

**Table1.** Network regions that significantly contribute to AMP, SLP, TEMP

| # | Main regions of the intrinsic brain network | AMP | SLP | TEMP |
| --- | --- | --- | --- | --- |
| 0 | Calcarine Cortex LR, SI LR | **-7.06\*** | **-6.72** | -4.28 |
| 1 | Somatosensory: SI, SII, ACC/PCC, and Thalamus R | **4.81** | -3.81 | 4.52 |
| 3 | Auditory Cortex LR | -3.38 | -4.55 | **5.43** |
| 5 | Somatosensory: SI LR, Cerebellum LR | 3.67 | -4.14 | **6.30** |
| 6 | Lateral Occipital Cortex LR | **5.67** | 3.27 | 4.20 |
| 7 | Precuneus, PCC, Lateral Occipital Cortex LR | -3.84 | -2.94 | **-5.33** |
| 10 | Medial Orbitofrontal Cortex LR, pACC L | **4.52** | 5.20 | **-6.20** |
| 11 | Lateral Occipital Cortex, Superior Frontal Gyrus L | **5.99** | **-6.70** | -2.90 |
| 12 | Lateral Occipital Cortex, Superior Frontal Gyrus R | **5.15** | 4.68 | 3.18 |
| 13 | Precentral Gyrus LR | **-4.17** | -2.95 | 2.96 |
| 14 | Cerebellum LR | **-4.36** | -2.96 | -3.41 |
| 18 | Lingual Gyrus, Fusiform Gyrus, Lateral Occipital Cortex LR | -3.64 | **6.16** | -3.99 |
| 20 | Frontal Paracingulate gyrus, Precuneus, Orbitofrontal Gyrus LR | **4.70** | -3.45 | -4.32 |
| 22 | Superior Frontal Gyrus, Lateral Occipital Cortex L | **4.29** | 5.02 | -5.06 |
| 23 | Supramarginal Gyrus L | 3.27 | **-5.47** | 4.03 |
| 29 | Inferior Frontal Gyrus, Superior Temporal Gyrus, Marginal Gyrus R | **5.64** | **6.14** | 3.62 |
| 32 | Middle Frontal Gyrus L | **4.39** | -4.68 | 4.65 |
| 34 | Frontal Pole, Insular Cortex L | **4.33** | -5.09 | 2.95 |
| 37 | Frontal Pole, Anterior Insula L | 2.47 | -4.12 | **6.02** |
| 38 | Frontal Pole L | **4.10** | -5.09 | **6.21** |
| 39 | Paracingulate Cortex, Medial Frontal Cortex, Accumbens LR | 3.66 | 4.72 | **-7.45** |
| 40 | Precuneus, Angular Gyrus LR | **-4.97** | 3.40 | -2.96 |
| 43 | ACC, Insular Cortex LR | **-4.27** | -3.81 | 4.56 |
| 44 | Insular Cortex, Paracingulate Cortex LR | -3.78 | **-5.47** | **6.18** |
| 55 | Middle & Superior Temporal Gyrus LR | -3.41 | 3.19 | **-7.85** |
| 57 | Frontal Pole R | -2.71 | -3.47 | **6.35** |
| 62 | Occipital Pole R | **7.41** | **-10.37** | **-5.35** |
| 66 | Occipital Pole L | **-7.39** | **6.77** | 3.48 |
| 67 | Thalamus, Brainstem LR | **-4.14** | 3.38 | -3.82 |
| 69 | Frontal Pole R, Cerebellum | **4.48** | -5.04 | 3.67 |
| 70 | Precentral Gyrus, Postcentral Gyrus, Inferior Frontal Gyrus LR, Anterior Insula R | **-4.09** | **-6.27** | **8.16** |
| 72 | Frontal Pole R | **-4.79** | **-5.53** | 4.38 |
| 78 | Frontal Pole LR, Superior Frontal Cortex L | **-4.43** | **-7.48** | **5.85** |
| 81 | Occipital Pole R | **11.10** | **8.92** | **6.91** |
| 94 | Superior Frontal Gyrus LR, Lateral Occipital Cortex R | **-5.07** | 5.08 | 4.31 |

\* **Bold numbers indicate significance (t-values; $p < 0.05$ , PALM corrected). Capital letters indicate the lateralisation of the components (L=left, R=right; LR=bilateral). From the total number of 100 components, the table includes only the components that are significant in either of the 3 entities: AMP, SLP, TEMP. The maps of all 100 components can be found in the Supplementary Material.**

### SLP - direction of pain intensity changes

For most networks that encode intensity changes, we found a negative relationship between increasing pain and network activity (Figure 4, Table 1). We found this effect for a network which predominantly includes the bilateral frontal pole and the left superior frontal cortex (#78). A further negatively-related network consists of the bilateral pre- and postcentral gyrus, the bilateral inferior frontal gyrus and the right anterior insula (#70). We found a local network in the left supramarginal gyrus (#23), and a larger network comprising the bilateral insular and paracingulate cortices to be negatively-related to the change of pain intensity direction (#44). Please note that the drop of brain activity occurs after the peak of pain intensity (#70, #78). There are further negatively-related networks predominantly in occipital areas (Table 1).

**Figure 4.**
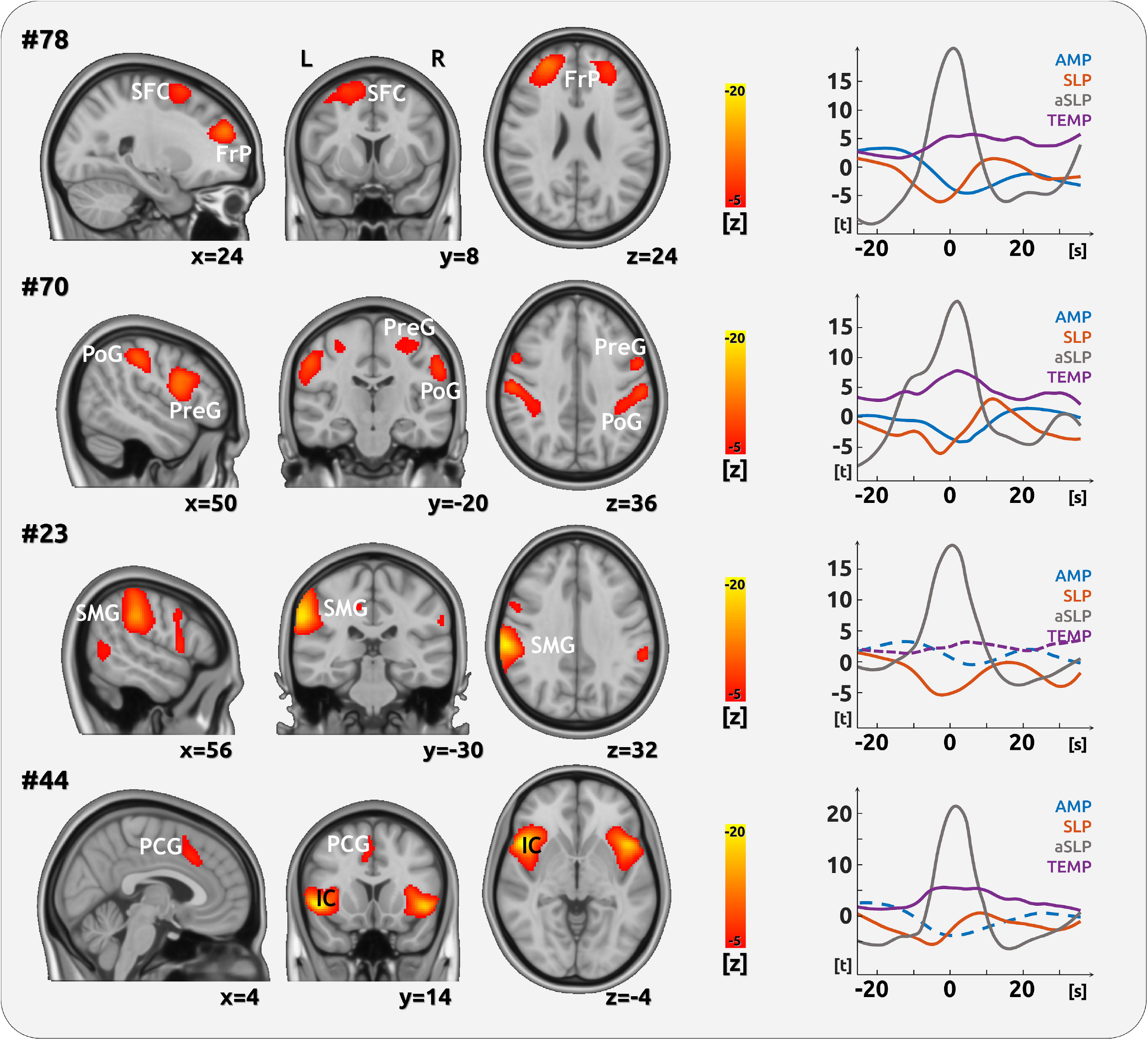
Encoding of the change of pain intensity ratings. The figure shows intrinsic functional networks that encode the intensity change (SLP) of tonic pain. The left side depicts the spatial characteristics of the networks; they were defined by the ICA. The significance of the contribution of each voxel to the network is represented by z-values. Only the right side depicts the pain intensity change encoding. The graph shows the timing of the network activity in reference to the current change of pain rating at time point 0. Solid lines indicate significant relationships, dashed lines indicate non-significant relationships. Please note the limited interpretability of the timing aspects due to the unknown lag of the haemodynamic response function, which may be different for each network. However, there is a *relative* delay for the change encoding compared to the pain intensity encoding for components #70 and #78. (Abbreviations: SFC superior frontal cortex, FrP frontal pole, PoG postcentral gyrus, PreG precentral gyrus, SMG supramarginal gyrus, PCG paracingulate gyrus, IC insula cortex).

A single fronto-temporal network (#29) that also encodes pain intensity is positively-correlated with the direction of pain intensity changes: rising pain is related to higher network activity (Figure 3). Please note the temporal lag between the SLP peak and the subsequent AMP peak.

### TEMP - encoding of small temperature fluctuations

We found a positive relationship between network activity and heat change for two bilateral somatosensory networks (#70, #5, Figure 4 and 5, Table 1) and two fronto-insular networks (#44, #37, Figure 4 and 5, Table 1). The representation of temperature change exhibits a complex picture as there are further networks consisting of frontal and occipital regions, which are positively-related to the heat pain changes (#81, #57, #38, #78, Table 1).

**Figure 5.**
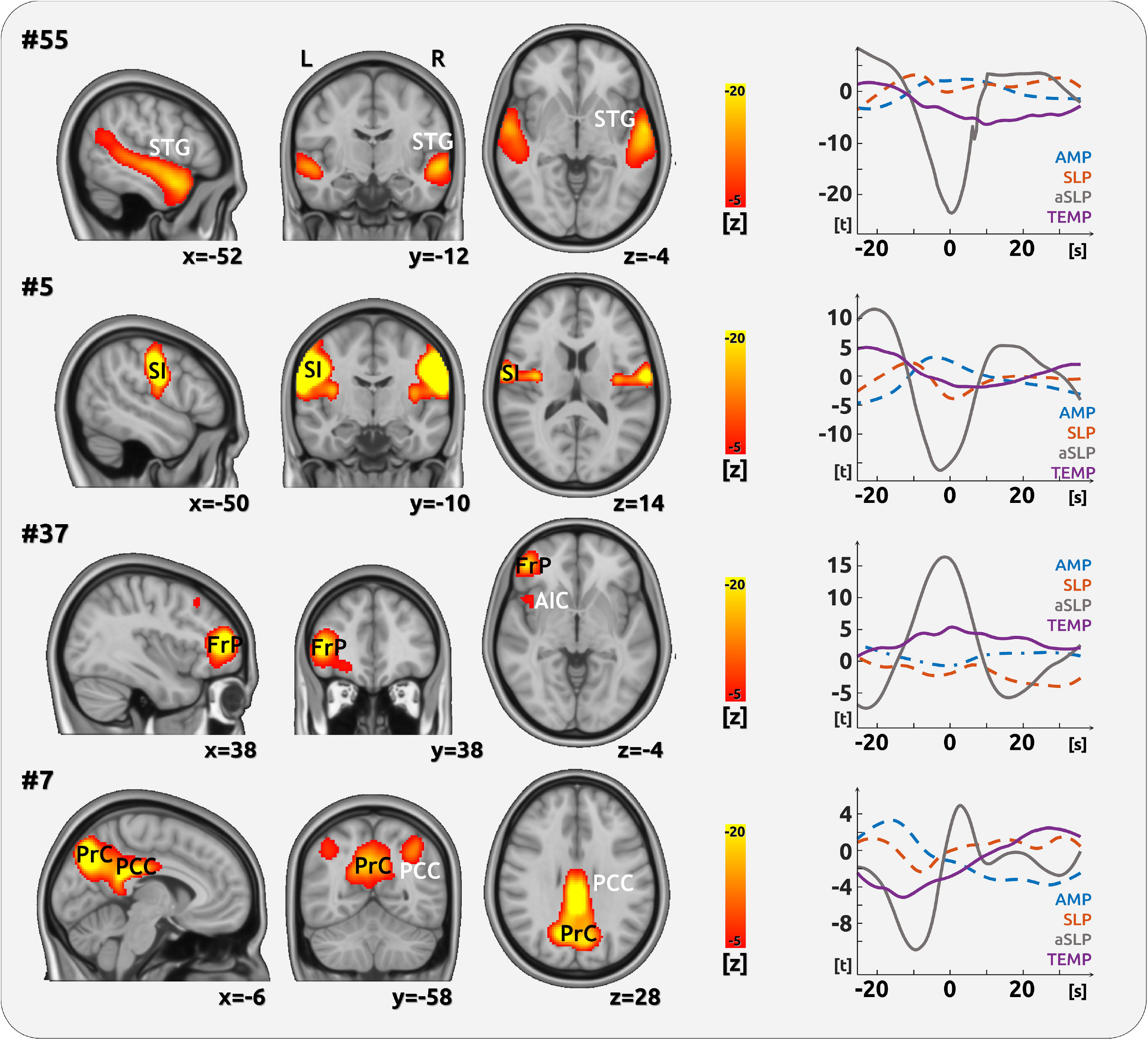
Encoding of the applied temperature. The figure shows intrinsic functional networks that encode the temperature change (TEMP) of tonic pain. The left side depicts the spatial characteristics of the networks; they were defined by the ICA. The significance of the contribution of each voxel to the network is represented by z-values. Only the right side depicts the temperature change encoding. The graph shows the timing of the network activity in reference to the current change of temperature at time point 0. Solid lines indicate significant relationships, dashed lines indicate non-significant relationships. Please note the limited interpretability of the timing aspects due to the unknown lag of the haemodynamic response function, which may be different for each network. Temperature change is also encoded in components #10 (Figure 3), #70, and #78 (Figure 4) (Abbreviations: STG superior temporal gyrus, SI primary somatosensory cortex, FrP frontal pole, AIC anterior insular cortex, PrC precuneous, PCC posterior cingulate cortex).

Negatively-related networks comprise a bilateral temporal network (Figure 5, #55), and a posterior network comprising the precuneous and the PCC (#7, Figure 5, all Table 1). Further networks that are negatively-related to temperature changes comprise the medial frontal cortex, frontal regions, the ACC, the paracingulate cortex, and the nucleus accumbens (#39, #10, table 1).

## Discussion

In the present study we explored how distinct aspects of the experience of tonic pain are reflected in the dynamically-evolving time course of intrinsic network activity. We were able to disentangle the contribution of a) the amplitude of pain intensity (AMP), b) the change of direction of pain intensity (SLP), and c) the fluctuations of temperature (TEMP) in relation to the spatiotemporal characteristics of the cortical networks. We could distinguish in two somatosensory networks (a) the cognitive-discriminative aspects that relate to the subjective perception of pain, and (b) the temperature-related aspects that are involved in the ascending spinothalamic stream of peripheral information transmission.

Language-related areas are suggested to play a supportive role during the evaluation process, particularly for rising and high pain states. A more extended discussion on methodological aspects can be found in Supplementary File 1.

### Disentangling distinct entities of tonic pain processing

#### AMP - pain intensity encoding

The amplitude of a somatosensory component (#1), consisting of SI, SII, the anterior part of the PCC, the posterior part of the ACC, and the thalamus, was found to encode pain; higher levels of the network activity were accompanied by higher levels of pain intensity. This finding is in line with previous studies emphasising the somatosensory-related cognitive-discriminative aspects of pain perception (Bushnell et al. 1999; Bingel et al. 2004).

In addition, we found a right-hemispheric network (#29) consisting of the inferior frontal, temporal, and marginal gyri. This network is suggested to fulfill an important role in the cognitive evaluation of painful stimulation, which may also be language-supported. For this component, there is a lag between the processing of pain change and the processing of pain intensity; the component is detecting and processing the increase in pain before it continues to process pain intensity. The contribution of temporal regions within the network corroborates recent findings on the involvement of the temporal cortex in pain processing in general (Atlas et al. 2010; Ayoub et al. 2019). Specifically, mnemonic aspects of pain unpleasantness encoding have also been related to temporal processing (Houde et al. 2020). Please note, this network is more active for higher pain than for lower pain, suggesting a stronger need for an in-depth evaluation of higher and more unpleasant pain intensities.

Further pain intensity encoding networks involve the contribution of various areas of the prefrontal cortex (particularly dorsolateral and medial areas), which are functionally connected to (a) the precuneous and the orbitofrontal cortex (#20), (b) the left posterior ACC (#10), (c) the cerebellum (#69), and (d) the left anterior insula cortex (#34). Regions within the frontal cortex were repeatedly shown to be involved in the cognitive and emotional modulation of phasic pain (Lorenz et al. 2003; Kalisch et al. 2006; Villemure and Bushnell 2009; Wiech 2016; Schulz et al. 2019).

We found a suppression of visual activity in occipital networks due to painful stimulation; this effect has been shown in previous studies (Bingel et al. 2007; Wager et al. 2013). The more lateralised occipital effects are likely to be caused by input from visual feedback; for higher pain intensities the cursor was located in the right visual hemifield, for lower pain intensities the cursor was located in the left visual hemifield.

The unknown delay of the HRF makes it impossible to interpret the timing of effects that are detected after or at the time point of the current pain rating. It is very likely that some of the effects may have been processed in the brain before the current rating; the sluggish BOLD response, however, has resulted in a delayed appearance of these effects. In contrast, effects that are found before the current rating can only have occurred and have been processed before the current rating. Consequently, for two components we found that preceding network activity determines the current experience of pain (#1, #10). This first (#1) component comprises somatosensory regions; these regions would be the first to receive incoming signals from the spinothalamic tract (Dum et al. 2009; Liang et al. 2011). Higher activity of a further component (#10) is associated with higher subjective pain levels despite lower levels of temperature. This component is mainly located in the orbitofrontal and anterior cingulate cortices that control the descending pain-modulatory pathways (Tracey and Mantyh 2007; Eippert et al. 2009); the combination of high pain and low temperature may reflect the attempts of the participants to activate pain inhibition in a low-temperature range. Activation of this system in high-pain phases may have been rendered unfruitful, had no effect on pain rating, and therefore no statistical outcome.

#### SLP - direction of pain intensity changes

An important dynamical aspect during the perception of fluctuating tonic pain is the processing of the direction of subjective pain intensity change. In our balanced design, we have an equal amount of phases with increasing and decreasing pain. Across all data, both exhibit a similar pain intensity, motor response, and decision-making, but differ in the direction the pain is progressing (up vs. down). One can assume that falling pain is more pleasant than rising pain (Baliki et al. 2010). Although the unknown delay of the HRF does not allow us to fully interpret the timing across networks, we can indeed cautiously consider the timing aspects for networks, where we observe an effect for both AMP and SLP (#29, #78, #70). Here, the cortical processing of the changes always precedes the processing of intensity, which is in line with logical reasoning.

For the processing of rising pain, in the right-sided intrinsic network that comprises the inferior frontal, the superior temporal, and the marginal gyri (#29), we can see an initial increase of network activity towards the time point of the current rating. The striking overlap with anterior and posterior language areas (Schulz et al. 2008, 2009) suggests this network integrates memory and inner speech, which appears to be particularly active for increasing pain and high pain (see above for the AMP results). We therefore suggest this network of brain regions is involved in inner verbalisation to support the rating process. Moreover, increasing pain lowers and potentially disrupts network activity. We found decreasing network activity towards the time point of the rising-pain in (a) a bilateral network consisting of the paracingulate and the anterior insular cortex (#44), (b) a supramarginal network (#23), (c) a bilateral sensory-motor network that also includes the right insula (#70), and (d) a bilateral frontal network (#78). All of these networks show lower activity with rising pain, the two latter at sustained high levels of pain. We can only speculate to which physiological or psychological processes are hampered here, however, the somatosensory (#70) and the insular components (#44) suggest that non-pain related aspects of higher-order (somato-)sensory integration are impaired during intense and rising phases of pain (Takeuchi et al. 2019).

#### TEMP - encoding of small temperature fluctuations

Due to the constantly-adapted temperature and the slow sensitisation-related decrease in temperature throughout the experiment, the temperature encoding can only reflect the relative and small changes of painful heat.

Indeed, the cortical processing of temperature changes is an important aspect of the current study as it represents the physical and objective aspects of the study. We found a number of networks that are positively-related to the magnitude of temperature changes. Some networks peak well before the evaluation of the current pain (at time point 0, see Figure 5). This confirms our observation during the recordings: there was a substantial lag of several seconds between the change of temperature and the conscious realisation of pain intensity change as indicated by a slider movement. Consequently, the cortical data exhibit two phenomena. *Firstly*, we found an early increase in a somatosensory network (#5). The activity in this network is unrelated to the subjective perception of pain intensity. *Secondly*, two networks which can be considered as the anterior (#10) and posterior default mode network (#7), exhibit an early decrease of cortical network activity. Although we can not precisely define the exact timing of the temperature-related processes, we can assume with certainty that these processes occur well before the current pain perception. This suggests a pre-conscious suppression of cortical processes, even before the participants were realising and rating the change in pain intensity. Unfortunately, the previous EEG studies on tonic pain perception did not take these timing effects into consideration (Schulz et al. 2015; Nickel, May, Tiemann, Postorino, et al. 2017). A further network that includes the nucleus accumbens (#39) is also connected to the physical characteristics rather than to the subjective perception of rising and falling pain (Baliki et al. 2010).

## Summary

We have investigated and disentangled the objective (temperature) and subjective (pain rating) aspects of the cortical processing of tonic pain. Through the dissociation of cortical processes into intrinsic networks, we have revealed distinct somatosensory processes that contribute to the encoding of pain intensity and temperature fluctuations. The cascade of subsequent processing steps demanded a systematic investigation of the cortical activity that flank the present moment of the evolving stream of conscious pain perception. Dynamic processes build up towards a current subjective percept of pain. Preconscious processing is related to changes of temperature and affects the default mode network.

## Data availability

The data that support the findings of this study are available from the corresponding author upon reasonable request.

## Acknowledgements

We thank Dr Stephanie Irving for copy-editing the manuscript.

