## Supplementary File 1 for "Intrinsic Network Activity Reflects the Fluctuating Experience of Tonic Pain"

### Challenges and advances of the study of intrinsic network activity during long-lasting pain perception

#### Controlling for sensitisation during prolonged painful stimulation

To investigate the encoding of long-lasting pain, it is paramount to keep the intensity of the subjective experience of pain fluctuating at a similar level across the time course of the experiment. Our current and previous data (Schulz et al. 2015) show that most of the participants exhibited sensitisation during the experiment, specifically requiring lower heat levels at the end of the experiment compared to the beginning. Ignoring the obligation to adapt the stimulation intensity will inevitably cause a systematic bias by introducing an effect of order (Favilla et al. 2014; Nickel et al. 2017). This is also true for the investigation of chronic pain, where an imbalanced rating time course can result in obtaining an effect in the first part of the experiment which may disappear in the second part due to a potential ceiling effect in some subjects (May et al. 2018).

#### A balanced design is a mandatory

Time series of functional imaging data require high-pass filtering; the fact that we are aiming to find a covariation of (a) the time series of behavioural data with (b) a filtered time series of the imaging data requires the same filtering for the behavioural data. For slowly rising or falling pain (Hashmi et al. 2013; Favilla et al. 2014; May et al. 2018), this would eliminate much of the variance in both pain ratings and imaging data. This results in analyses where the most essential effect (brain ~ pain rating) can not be computed, ultimately leading to analysis of the effects of rising versus stable phases, which includes motor and decision-making activity for the rising part but not for the stable part (Hashmi et al. 2013). The major advance in the current balanced stimulation is that we can investigate the intensity of pain without an effect of order, as well as the cortical underpinnings of rising and falling pain without any systematic motor or decision-making confound.

#### Functional dissociation of AMP, SLP, aSLP

Due to the balanced design we were able to dissociate three entities of the paradigm: AMP, SLP, aSLP. The encoding of AMP is independent from rising pain as its computation is based on phases of rising pain as well as phases of falling pain. In a similar vein, motor activity, decision making occur at all levels of AMP and are equally frequent during rising and falling pain. Please note that falling pain is not synonymous to low pain. Therefore, AMP and SLP are not interchangeable concepts. AMP is based on periods of rising and falling pain. Both exhibit different temporal characteristics (see time courses in Figures 3, 4, and 5).

#### Dynamics of intrinsic network activity: peaks and troughs

The current within-subject approach is unprecedented in neuroimaging. The analysis of the trajectory of ongoing intrinsic networks is largely applied to investigate the connectivity between networks (Smith et al. 2015; Schumacher et al. 2018). In pain research, previous studies were predominantly pursuing analyses on non-stationary aspects, and compared chronic pain patients with healthy controls regarding 3D maps of certain components like the default mode network (Napadow et al. 2010; Loggia et al. 2013; Baliki et al. 2014) or the salience network (Kucyi et al. 2013; Kim et al. 2018; van Ettinger-Veenstra et al. 2019). These maps are interpreted as representing the *stationary* strengths of functional connectivity across the entire recording period (Buckner et al. 2013). By merely focusing on cortical maps leaves open the question of whether there are different physiological or cognitive states during peaks and troughs of the network time series.

##### Dynamics of intrinsic network activity: temperature encoding

It must be noted for the current study that we can not investigate the cortical underpinnings of the absolute amplitude of temperature with a standard EPI sequence; Figure 1 shows a gradient of temperature decline across the time course of the experiment. This gradient has three implications. *Firstly*, by using an unfiltered temperature gradient, we would introduce an effect of order where we would contrast the first part with the last part of the experiment. *Secondly*, a substantial share of the temperature gradient is likely related to peripheral sensitisation and can not be disentangled from cortical sensitisation. *Thirdly*, the sensitisation occurs at a very low frequency across the time course of the experiment. Such low-frequency aspects are inevitably filtered out from the cortical data. More recent developments in imaging techniques, such as ASL (Segerdahl et al. 2012, 2015), would be able to capture low-frequency gradual changes in the data but would also need to take into account the potential effects of order.

Nevertheless, the filtered data allows us to investigate how the small and high-frequent changes of temperature are subserved in the human brain. The findings corroborate a remarkable dissociation between the processing of objective (temperature) as compared to subjective (pain ratings) measures of pain (Schulz et al. 2015).
